# Cellular contributions to ictal population signals

**DOI:** 10.1101/2022.07.08.499193

**Authors:** Lauren A. Lau, Zhuoyang Zhao, Stephen N. Gomperts, Kevin J. Staley, Kyle P. Lillis

## Abstract

**Objective:** The amplitude of ictal activity is a defining feature of epileptic seizures, but the determinants of this amplitude have not been identified. Clinically, ictal amplitudes are measured electrographically (using e.g. EEG, ECoG, and depth electrodes), but these methods do not enable the assessment of the activity of individual neurons. To identify the cellular determinants of the ictal signal, we measured single cell and population electrical activity and neuronal calcium levels via optical imaging of the genetically encoded calcium indicator (GECI) GCaMP.

**Methods:** Spontaneous seizure activity was assessed in an awake, behaving mouse model of focal cortical injury and in organotypic hippocampal slice cultures (OHSC), an in vitro preparation from which recurrent seizures can be readily captured. Single cell calcium signals were linked to a range of electrical activities by performing simultaneous GECI-based calcium imaging and whole-cell patch-clamp recordings in spontaneously seizing OHSCs. Neuronal resolution calcium imaging was then performed during spontaneous seizures in vivo and in vitro to quantify the cellular contributions to the population-level calcium signal.

**Results:** Population signal may increase from three potential sources: 1) increased synchrony, i.e. more co-active neurons, 2) altered active state, from bursts of action potentials and/or paroxysmal depolarizing shifts in membrane potential, and 3) altered subthreshold state, which includes all lower levels of activity. The largest contributor to the signal recorded at seizure onset was increased subthreshold activity, consistent with either barrages of excitatory postsynaptic potentials or sustained membrane depolarization. The relative contribution of synchrony increased as seizures progressed, but cell intrinsic alterations in both the subthreshold and active states remained the largest driver of the ictal signal.

**Significance:** We introduce here a novel method for the quantification of the relative contributions of inter-versus intra-cellular changes to provide a critical link between single neuron activity and population measures of seizure activity.

**Key Points:**

- Neuronal calcium as measured by GCaMP reports a range of membrane depolarizations, from EPSPs to action potential firing and paroxysmal depolarizing shifts
- The mean population calcium signal is highly correlated with the electrographic local field potential
- Increased calcium signal during seizure onset is largely driven by increased subthreshold calcium within individual neurons
- Recruitment of newly active neurons is a minor contributor to the increasing population-level signal during the transition to frank seizure

## INTRODUCTION

The amplitude of paroxysmal ictal activity is a defining feature of epileptic seizures. Although low-amplitude seizures occur, for example in infantile spasms (*1*), this is the exception. The rule is that seizure activity is recognizable in electroencephalographic recordings in large part because of the distinguishing increase in EEG amplitude (*2, 3*). However, despite the central role of EEG in diagnosing epilepsy and localizing seizure foci for surgical planning, the determinants of this unique increase in amplitude remain unclear. Here we identify those determinants.

The EEG signal is a nonlinear summation of the activity of the neurons and glia in the underlying cortex (*4, 5*). Micro-electrode array-based electrophysiological methods have been used to record the activity of individual neurons with high temporal and moderate spatial resolution during physiological activity and, in some cases, even during seizure onset (*6, 7*). However, these methods are limited to capturing the spiking activity of neurons. As action potentials are too brief to be reflected in EEG, the extracellular population signal is largely attributed to post-synaptic potentials. An additional contributor to ictal signal is paroxysmal depolarizing shifts (PDS) in the neuronal membrane potential, which lasts hundreds of milliseconds (*8, 9*). As EEG reports a spatial average of underlying cortical activity, only activity that is synchronous across the reported cortical volume is recognizable(*10*). Epileptiform EEG has therefore been considered to represent highly synchronous activity of cortical neurons (*5, 11*). Increased synchrony can arise from recurrent activation of neurons, or the recruitment of new populations of neurons (*7*).

To capture the relevant activities of individual neurons, we combined whole-cell recordings of the membrane potential and field potential recordings with optical imaging of calcium using the genetically encoded calcium indicator GCaMP. Small fluctuations in calcium accompany postsynaptic potentials, as calcium enters through ionotropic glutamate receptors and certain classes of voltage-gated calcium channels (*12–14*). Neuronal spiking and PDS both drive high calcium signal (*15, 16*). In addition to these cell-intrinsic signals, network synchrony can also be measured in a straightforward way with calcium imaging by quantifying the number of co-active neurons. Thus, recordings with high spatial resolution and sensitivity to a wide dynamic range of signals can be obtained over a large cortical volume.

Simultaneous microendoscope calcium imaging and local field potential (LFP) recordings are now feasible in awake, behaving murine models of focal cortical injury (*17*) (where seizures were observed following cortical aspiration in the seizure susceptible APP/PS1 mouse model of early-onset Alzheimer’s (*18*)). However, spontaneous seizures while the subject is being recorded are exceptionally rare events, and only a single partial spontaneous ictal recording has been reported to date (*19*). The current report is based on the serendipitous recordings of several such spontaneous seizures *in vivo*. To extend these rare in vivo findings, we also performed multimodal recordings in a reduced, chronically epileptic preparation: the organotypic hippocampal slice culture (OHSC). Unlike in vivo models, seizures in OHSC are frequent and straightforward to capture. Additionally, cellular resolution recordings can be obtained from the entire epileptic network and contributions to ictal signal can be unambiguously ascribed to the seizure focus (*20*).

We developed novel methods to quantify the relative contribution of inter-cellular (i.e. synchronization) and intra-cellular sources to the overall ictal signal. During seizure onset the majority of the signal increase was driven by increased amplitude of subthreshold discharges of individual neurons, with the role of active state neurons becoming more prominent during the transition to frank seizure. We further show that synchronous activity during seizure is dominated by reactivation of neurons, rather than the recruitment of new populations of neurons into the seizure.

## METHODS

### Study approval

All animal protocols were approved by the Massachusetts General Hospital Subcommittee on Research and Animal Care and were conducted in accordance with the United States Public Health Service’s Policy on Humane Care and Use of Laboratory Animals.

### In vivo calcium imaging and electrophysiology

Virus injection of AVV1-hSyn-GCaMP6f and endoscope and GRIN lens implantation surgery was performed as previously described (*21, 22*), in an adult APP/PS1 mouse. Simultaneous calcium imaging (20 Hz) and electrophysiology recordings from a custom-made probe of 7 stereotrodes were performed. Additional details in Supplemental Methods.

### Organotypic Hippocampal Slice Culture (OHSC) calcium imaging and electrophysiology

OHSCs were prepared as described previously (*20*). OHSC generate spontaneous recurrent seizure-like events, henceforth referred to as seizures. Whole-cell patch-clamp recordings were performed with simultaneous hSyn-soma-GCaMP8m two-photon-based calcium imaging (30 Hz). Chronic hsyn-GCaMP7f calcium imaging (35 Hz) was performed in a custom imaging system, in combination with LFP recordings. Additional details in Supplemental Methods.

### Processing of calcium signaling

Neuronal regions of interest (ROI) were identified with the Fiji plugin TrackMate (*23*). The calcium signal of neurons was separated into two states: subthreshold and active. We developed an automated method for separating these states, by defining a detection threshold value calculated from a period of baseline inter-ictal activity. The baseline period was chosen to exclude high amplitude transients (characterized by a rapid increase in calcium amplitude). The normal variance in low amplitude calcium was then calculated to define a detection threshold which could reliable separate subthreshold and active amplitude activity. The mean population calcium signal (mean Δf/f) reflects changes in 3 variables: the fraction of synchronously active neurons, the mean active state calcium amplitude, and the mean subthreshold calcium amplitude. All analyses were performed in ImageJ (Fiji) and Matlab. Additional details in Supplemental Methods.

## RESULTS

### Genetically encoded calcium indicators (GECIs) to simultaneously record single cell and population seizure activity

To demonstrate the relationship between the calcium signal and epileptiform electrical activity at the single neuron level, we performed simultaneous whole-cell patch clamp recordings and GCaMP-based calcium imaging in the OHSC model (Figure 1),. To quantify when neurons are active (based on the calcium signal alone), a detection threshold was employed to separate subthreshold and active states(see Methods for details). During inter-ictal periods, the subthreshold state is characterized by a relatively stable, low amplitude calcium signal. (Figure 1, n = 3 CA1 pyramidal cells from 3 OHSCs) and was characterized by a low frequency (<1 Hz) of excitatory postsynaptic potentials (EPSPs). Sparse action potential firing may also occur in the subthreshold calcium state (*16, 24*).

**Fig. 1.**
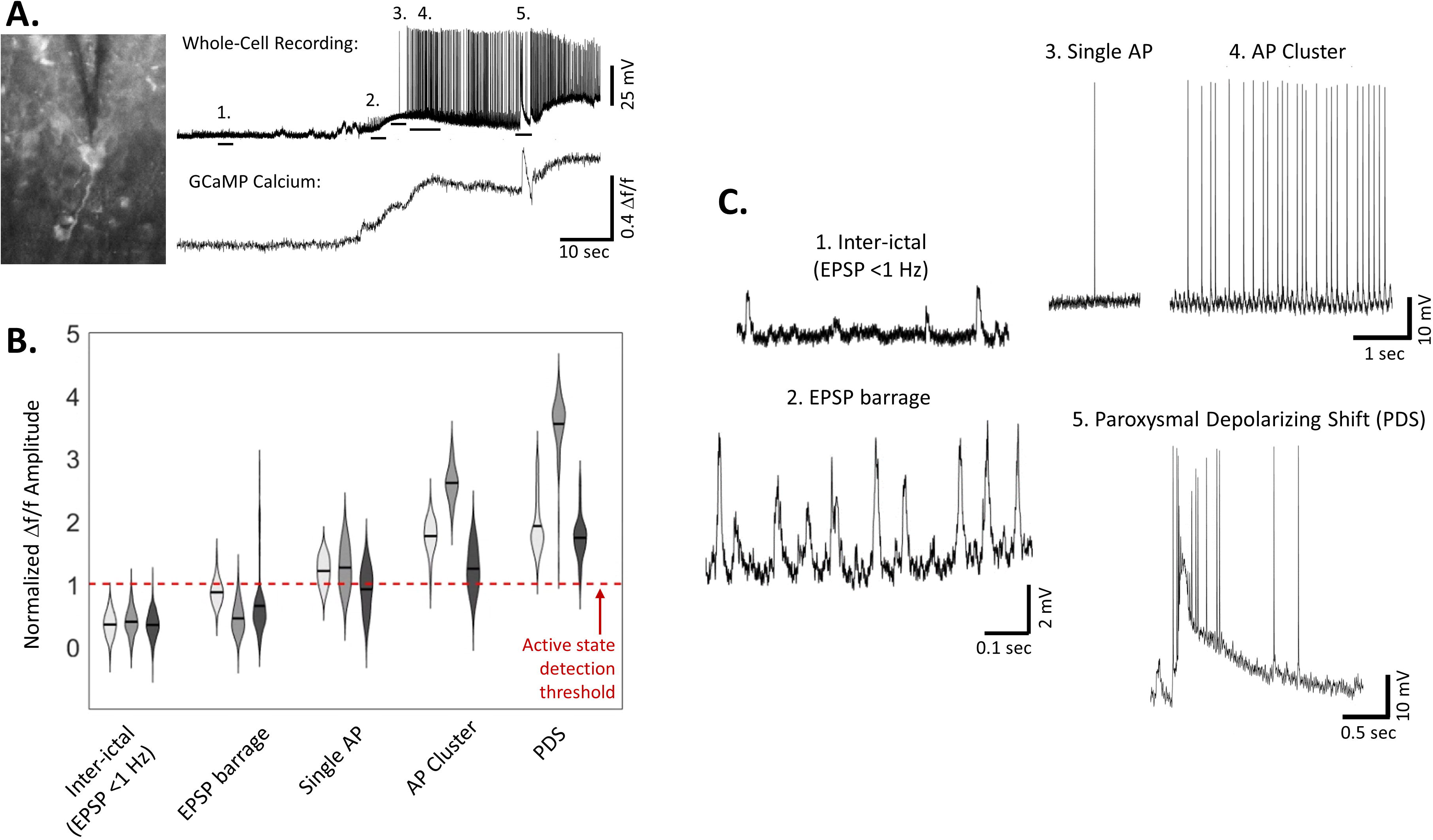
Paired calcium imaging and whole-cell patch clamp. (**A**) Left: Image of GCaMP+ neuron and patch pipette in OHSC. Right: Paired current-clamp recording and soma-GCaMP8m calcium signal (Δf/f) during spontaneous seizure. (**B**) Violin plot of calcium Δf/f amplitudes during different types of electrical activity, normalized to active state detection threshold per cell. (n = 3 pyramidal cells, from 3 OHSCs). (**C**) Zoom-in to each category of electrical activity, as noted in A. 1: Inter-ictal activity is defined as period between seizures where EPSPs frequency is below 1 Hz; 2: EPSP barrage, typically seen at seizure onset, with EPSP frequency >1 Hz; 3: Single Action Potential (AP), defined as only AP in a 150 ms window. 4: AP Cluster, defined as multiple (10+) action potentials with <150 ms interval between spikes; 5: Paroxysmal depolarizing shift (PDS), characterized by action potentials on top of a depolarized plateau of 20-50 mV, lasting tens-hundreds of ms.

During seizure onset, we observed that a barrage of EPSPs (frequency >4 Hz) is capable of driving an increase in the amplitude of the calcium signal (Figure 1), which still remained subthreshold. Isolated action potentials (inter-spike interval > 150 ms) were frequently observed during seizure onset and resulted in higher amplitude, active state calcium signal. Clusters of action potentials (defined as 10+ APs with inter-spike interval <150 ms) observed during spontaneous seizures drove the calcium amplitude even higher. Paroxysmal depolarizing shifts were associated with the highest amplitude calcium signal (Figure 1). Interneurons were not typically observed with PDS but often displayed high frequency firing during spontaneous seizures, which consistently corresponded to an active state calcium response (n = 3 interneurons, Figure S1).

The network calcium signal can easily be obtained by averaging the calcium signal of all neurons in the field of view. **Seizures** were defined from the population-level calcium signal as periods of high amplitude calcium events lasting >10 seconds that coincided with increased amplitude in at least 1 channel of the LFP (Figure 2C-F). **Seizure onset** was defined as the period of relatively lower amplitude activity when the population calcium could first be distinguished from inter-ictal activity until the transition to **frank seizure**-marked by a local maximum in the population calcium slope (Figure 2E and F). Frank seizure is also referred to as **ictal** activity.

**Fig. 2.**
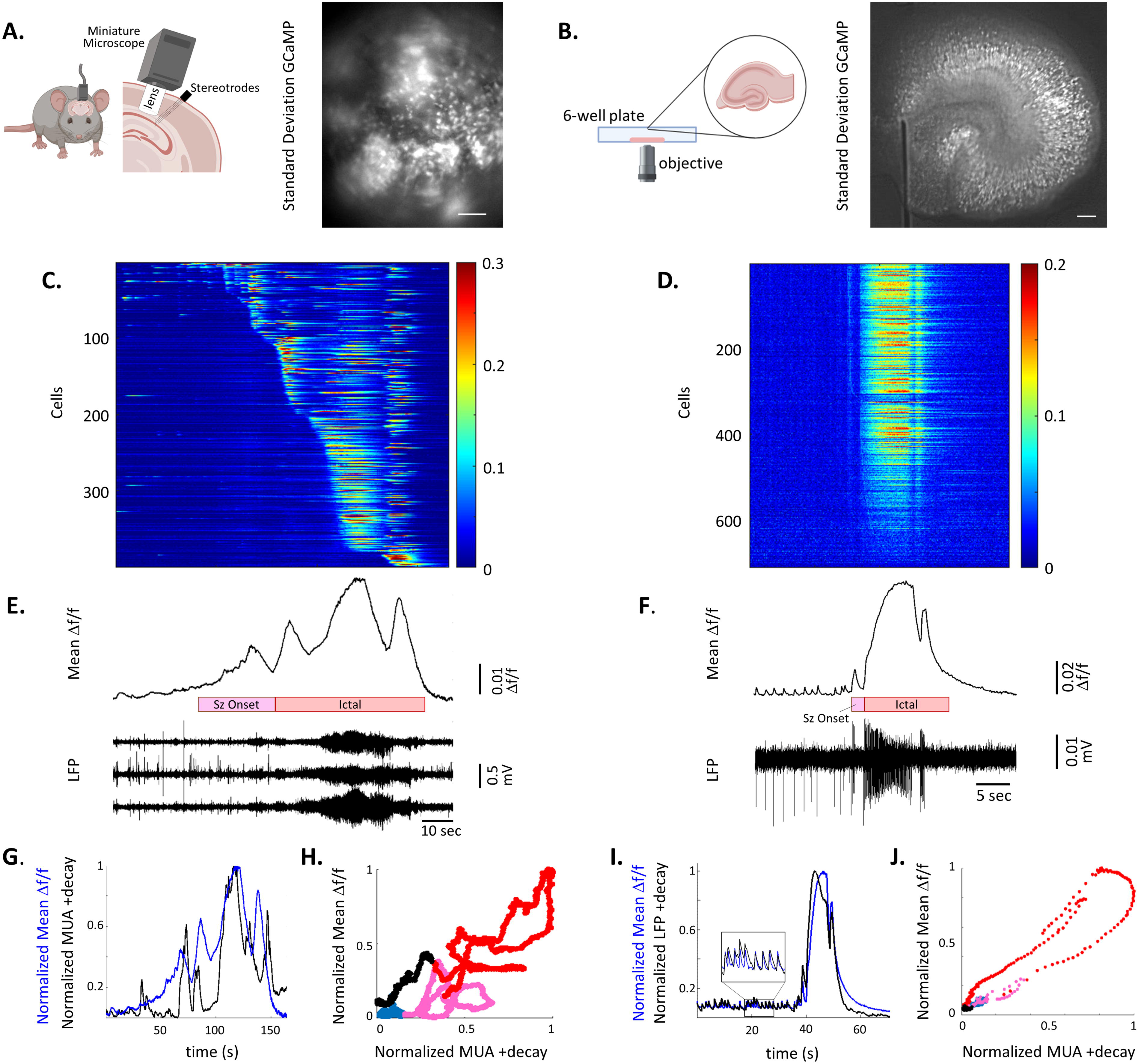
Paired calcium imaging and local field potential in vivo and in vitro. (**A**) Left: schematic of in vivo imaging. Right: Representative image of standard deviation projection during syn-GCaMP6f calcium imaging. Scale bar = 100 μm. (**B**) Left: Schematic of in vitro imaging. Right: Representative image of standard deviation projection during syn-GCaMP7f calcium imaging. Scale bar = 100 μm. (**C**) Raster plot of GCaMP6f Δf/f of individual neurons during in vivo seizure (**D**) Raster plot of GCaMP7f Δf/f of individual neurons during in vitro seizure (**E**) Paired mean GCaMP6f Δf/f trace and 3 channels of local field potential (LFP), in vivo. (**F**) Paired mean GCaMP7f Δf/f trace and LFP, in vitro. (**G**) Plot of normalized local field potential activity that has been high-pass filtered at 300 Hz for multiunit activity (MUA), rectified and down-sampled to rate of imaging (black) and normalized mean Δf/f calcium (blue), simultaneously recorded during in vivo seizure (**H**) Correlation in signal between the two modalities, color indicates phase of seizure (blue = baseline inter-ictal; pink = seizure onset; red = ictal; black = post-ictal). (**I**) Plot of normalized local field potential activity that has been rectified and down-sampled to rate of imaging (black) and normalized mean Δf/f calcium (blue), simultaneously recorded during in vitro seizure (**J**) Correlation in signal between the two modalities, color indicates phase of seizure (blue = baseline inter-ictal; pink = seizure onset; red = ictal; black = post-ictal).

We recorded neuronal-resolution calcium activity both in vivo and in vitro. The in vivo recordings describe spontaneous seizures in awake, behaving APP1/PS1 mouse that had a cortical injury due to cortical aspiration/GRIN lens implantation (Figure 2A). Endoscopic imaging of neuronal hSyn-GCaMP6f calcium activity was paired with recording of the local field potential (LFP) and unit activity from 7 stereotrodes sampling the nearby cortical and hippocampal areas (as detailed in (*21, 22*)).

Capturing *spontaneous*, non-convulsive seizures in vivo is rare. Furthermore, the limited field of view in vivo does not allow for definitive identification of the seizure onset zone. We therefore also performed hSyn-GCaMP7f based calcium imaging in vitro, in the OHSC model (Figure 2B). Due to the reduced size of the OHSC, the entire network could be sampled, guaranteeing that we captured activity from the seizure focus (versus strictly propagated activity, as is likely to be the case in vivo). In both models, the somatic calcium signal from hundreds of individual neurons was extracted (Figure 2C-D), and calcium signals were corrected for motion, neuropil contamination and normalized as Δf/f (see Supplemental Methods).

We found a high correlation between the optical recordings of neuronal calcium activity and the electrophysiological recordings of population activity. To compare the two modalities in vivo, we transformed the LFP by filtering for multiunit activity (MUA), rectifying, down-sampling (to match the imaging rate), and convolving with GCaMP6f decay kinetics (*16*). We found a correlation of 0.7-0.84 between the mean population calcium activity and MUA activity, with the highest correlation occurring in the LFP channel closest to the site of imaging (Figure 2G and H). Rarely, in vivo seizures were observed where the increased amplitude in calcium was apparent tens of seconds earlier than the increased amplitude in the LFP, likely due to the imaged cells falling outside of the electrical field detected by the LFP electrode (Figure S2). These were analyzed as a separate class of seizures (Figure S3).

The LFP data was similarly transformed in vitro(*25*), revealing a correlation of 0.96 between the calcium signal and electrophysiology (Figure 2I and J). The comparatively higher correlation between population calcium and the LFP observed in vitro is likely due to the complete sampling of all neurons in the network (whereas neurons outside of the imaging field of view are likely to be contributing to the electrical activity in vivo).

### Calcium imaging to dissect population activity

Calcium imaging can thus be used to make inferences about several types of neuronal activity. Single cell calcium signals can be averaged in a straightforward manner to calculate the network calcium signal (Figure 3A-B). The fraction of active neurons was used to assess network **synchrony** (Figure 3C-E). Synchrony could be further separated into **recruitment** (i.e. new activation) and reactivation. Recruitment was based on each seizure event (from seizure onset to the end of the ictal period) such that, the first time a neuron was active in a seizure epoch was considered the recruitment point. Each neuron could only be recruited once per seizure, and activity prior to seizure onset did not inform recruitment. **Reactivation** refers to any time a neuron is in the active state after the initial recruitment point and includes both continued activity of a neuron after recruitment or renewed activation after a period of inactivity. In addition to changes in network synchrony, the amplitudes of the calcium signal in both the subthreshold and active states vary, consistent with differing modes of activity (Figure 3F). Together the changes in intra-cellular calcium amplitude, recruitment, and reactivation underly the population level ictal signal (Figure 3).

**Fig. 3.**
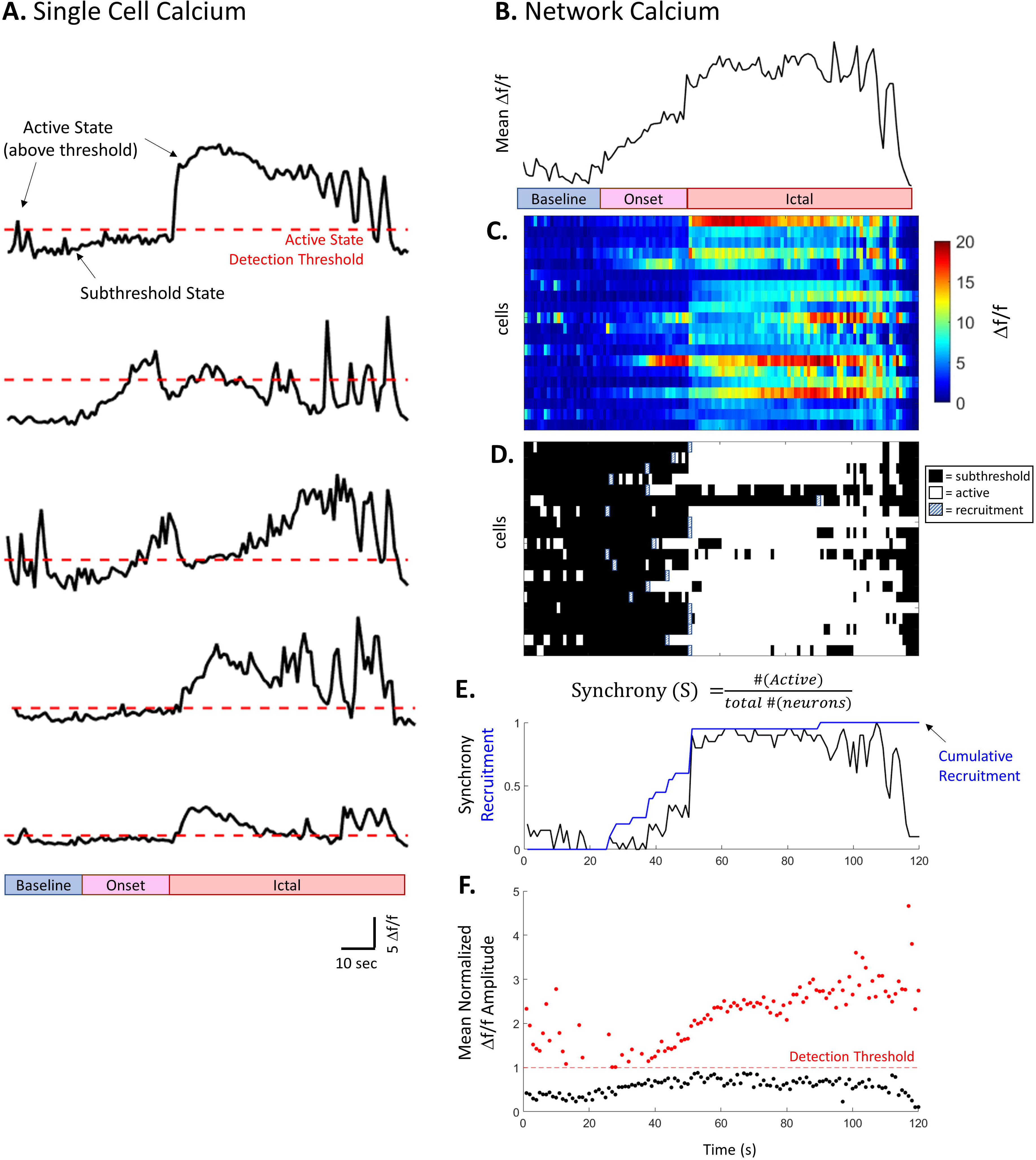
Changes in network synchrony and calcium amplitude underly population level changes. (**A**) Example single cell calcium traces. Red line indicates active state detection threshold for each cell. Phase of seizure indicated by colorbar. (**B**) Top: Mean Δf/f calcium of 20 neurons. (**C**) Raster plot showing the Δf/f amplitude for each of the 20 neurons. Neurons in (A) correspond to cell #1, 5, 10, 15 and 20 in raster plot. (**D**) Raster plot depicting when neurons are in the subthreshold (black) or active states (white). Recruitment shown in hatched white/blue and represents the first time a neuron is in the active state during the seizure epoch. (**E**) Plot of network synchrony over time (i.e. fraction of network in the active state). Blue line = cumulative recruitment. (**F**) Scatter plot of the mean active (red) and subthreshold (black) Ca^+2^ Δf/f amplitude over time normalized to detection threshold (red dotted line). Number of cells contributing to mean varies over time depending on number of neurons in active versus subthreshold state at each time point.

### Network synchrony during seizure

To quantify synchronous activity, we used the instantaneous fraction of active neurons (where fraction synchronously active (S) = # active neurons / total # of GCaMP-positive neurons, in each time frame of either 28 ms (OHSC) or 50 ms (in vivo)). During interictal activity, the mean fraction that were synchronously active was very low (0.2% ± 0.003, n = 82 recordings OHSC; 0.49% ± 0.08, n = 3 recordings in vivo). This interictal sample ranged from 11-56 seconds, depending on how much pre-seizure activity was captured. Across the same time period, 47.0% ± 0.6 (OHSC) and 7.2% ± 2.2 (in vivo) of neurons are active (above detection threshold) at least once, revealing low levels of asynchronous activity inter-ictally. The time-averaged fraction of synchronously active cells increased during seizure onset (Figure 3E and 4A-B, S= 6.7% ± 0.6, n = 82 seizures in OHSC; Figure 4F-G, S= 8.2% ± 2.2, n = 5 seizures in vivo). Increased synchrony was even more pronounced during ictal activity (S = 47.1% ± 1.9, in OHSC; S = 20.0% ± 4.2 in vivo). The percent active at each time point is calculated independently, and different neurons contribute to network synchrony at each time point.

**Fig. 4.**
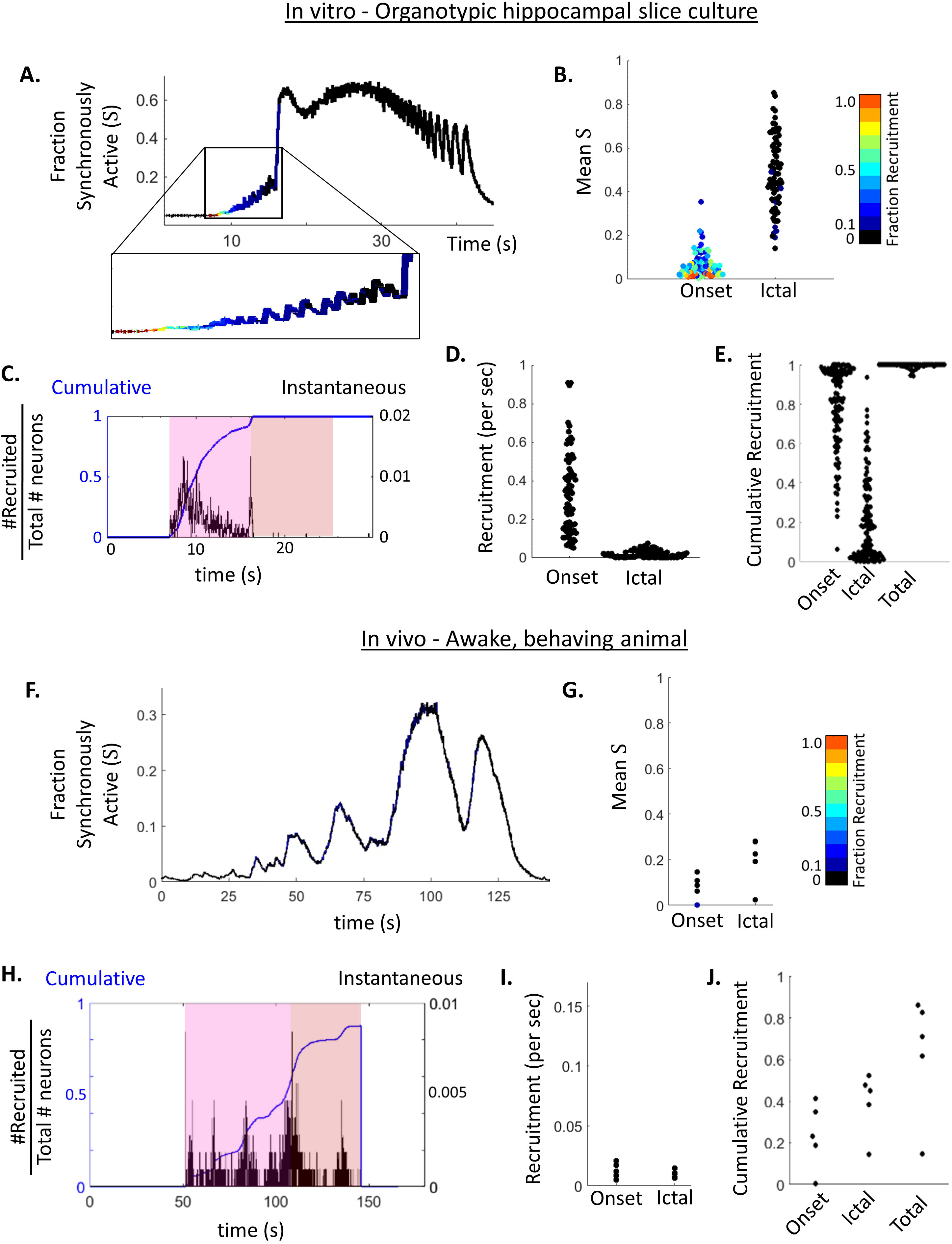
Network synchrony and recruitment. (**A**) Example plot of fraction of synchronously active neurons (S = # active neurons/total # neurons) over time from seizure in OHSC. Color represents fraction of S that is new recruitment. (**B**) Beeswarm plot of time-averaged fraction of synchronously active neurons (S) during seizure onset and ictal activity. Colorbar = recruitment/total fraction active. n = 82 seizures from OHSCs. (**C**) Plot of instantaneous recruitment (black) and cumulative recruitment (blue), from same seizure as (A). (**D**) Beeswarm plot of mean rate of recruitment (fraction of total network recruited per second) during onset and ictal periods, n = 82 seizures from OHSCs. (**E**) Beeswarm plot of cumulative recruitment during onset, ictal and in total (onset + ictal), n = 82 seizures from OHSCs. (**F**) Example plot of fraction of synchronously active neurons over time from an in vivo seizure. Color represents fraction of S that is new recruitment. (**G**) Beeswarm plot of time-averaged fraction of synchronously active neurons (S) during seizure onset and ictal activity. Colorbar = recruitment/total fraction active. N = 5 in vivo seizures. (**H**) Plot of instantaneous recruitment (black) and cumulative recruitment (blue), from same seizure in (F). (**I**) Beeswarm plot of mean rate of recruitment (fraction of total network recruited per second) during onset and ictal periods, n = 5 in vivo seizures. (**J**) Beeswarm plot of cumulative recruitment during onset, ictal and in total, n = 5 in vivo seizures.

New recruitment accounted for 32.2% ± 2.4 and 0.4% ± 0.2 of the total active fractions during onset and frank seizure in OHSC, (Figure 4A-C). In vivo, new recruitment constitutes 2.7% ± 1.8 and 0.4% ± 0.2 of the total active fractions during onset and frank seizure activity, (Figure 4F-H). The rate of new recruitment in OHSC was 32.5% ± 2.3 of total network recruited per second during onset and 2.0% ± 0.2 during frank seizure (Fig 4D); recruitment in vivo was much slower and occurred at a rate of 1.3% ± 0.01 (onset) and 1.0% ± 0.01 (ictal) (Figure 4I). In total, 78.6% ± 2.0 of neurons were recruited during seizure onset in OHSC, with the remaining 21.0% ± 2.0 of the neuronal population recruited during ictal activity, for a total cumulative recruitment of 99.6% ± 0.1 (Figure 4E). This reveals that the transition to frank seizure is largely due to synchronous activation of previously recruited neuronal populations in OHSC. In vivo, 23.7% ± 6.3 of neurons were recruited during seizure onset and 39.5% ± 6.0 during the ictal period, for a total cumulative recruitment of 63.2% ± 11.5 (Figure 4J). In vivo, while synchronous activity is predominately driven by reactivation, a significant number of neurons are being newly recruited during seizure progression. These differences in recruitment are likely due to differences in the models (where there are far more neurons available to recruit in vivo versus in vitro, and OHSC have a higher fraction of recurrent connections). But the differences may also reflect both slower and less total recruitment occurring outside of the seizure focus, as in vitro recordings were always in the seizure focus, but in vivo recordings may be either in the focus or the propagation area.

### Dynamics of the intra-cellular calcium amplitude

To quantify the change in intra-cellular calcium amplitude during seizure, we first identified a baseline period of non-ictal activity (without seizure or epileptiform activity). From this period, we calculated the mean baseline subthreshold (ST) calcium amplitude (STCa^+2^_t=0_) for each neuron by averaging all subthreshold values during the baseline window. We then normalized the subthreshold calcium amplitude 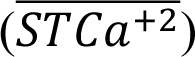 by dividing by STCa^+2^_t=0_ (Figure 5C), which provides a measure of the fold increase in subthreshold calcium amplitude over time. On average, the mean fold increase of the subthreshold calcium amplitude during seizure onset was 1.36 ± 0.02 (n = 82 seizures) in OHSC, and 1.68 ± 0.21 (n = 5 seizures) in vivo (Figure 5E and F). During frank seizure, the average increase in STCa^+2^ amplitude was 2.03 ± 0.02 in OHSC, and 1.80 ± 0.42 in vivo (Figure 5E-F). While, by definition, neurons are in the subthreshold calcium state at all points when they are included in the STCa^+2^ population, on average the subthreshold calcium amplitude is significantly increased during seizure onset (p < 0.001 OHSC; p = 0.01 in vivo, paired t-test) and frank seizure (p < 0.001 OHSC; p = 0.01 in vivo). To address the possibility of residual neuropil contamination driving this increase, we hand-drew regions of interest in a subset of neurons in OHSC (Figure S4A). In this case, nearly complete neuropil subtraction can be performed (see Supplemental Methods) and the increased subthreshold amplitude persists (Figure S4B; p = 0.003 (onset) and p < 0.001 (ictal). We also performed cellular resolution two-photon imaging of calcium with soma-targeted GCaMP8m during seizure in OHSC. These conditions greatly reduce the contribution of neuropil contamination, and the significant increase in subthreshold calcium amplitude is still observed (Figure S4C; p < 0.001 for onset and ictal).

**Fig. 5.**
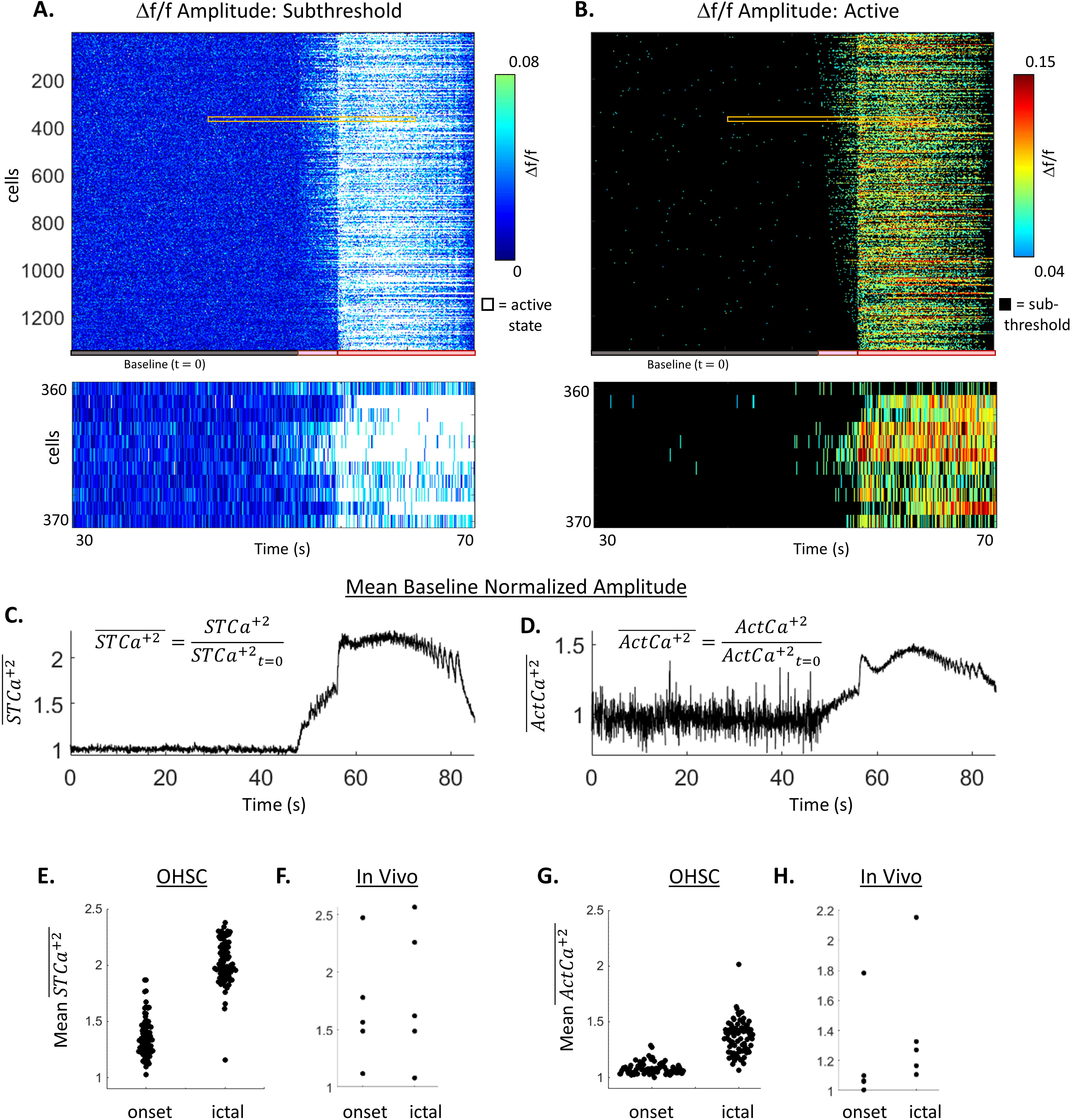
Change in calcium amplitude. (**A**) Raster plot of Δf/f signal of neurons in the subthreshold state. White = time points when the neuron was in the active state. Colors on bottom show stages of activity, gray = baseline activity, pink = seizure onset, red = ictal. Bottom: zoom-in to 40 seconds of activity in 10 cells. (**B**) Raster plot of Δf/f signal of neurons in the active state. Black = time points when the neuron was subthreshold. Bottom: zoom-in to 40 seconds of activity in 10 cells. (**C**) Example mean baseline normalized subthreshold calcium (STCa^+2^) amplitude over time of all neurons shown in (A). (**D**) Example mean baseline normalized active calcium (ActCa^+2^) amplitude over time of all neurons shown in (B). Data from A-D all show the same seizure recorded from a OHSC. (**E**) Beeswarm plot of time-averaged mean normalized STCa^+2^ amplitude during seizure onset and ictal in OHSC. n = 82 seizures. (**F**) Beeswarm plot of time-averaged mean normalized STCa^+2^ amplitude during seizure onset and ictal in vivo. n = 5 seizures. (**G**) Beeswarm plot of time-averaged mean normalized ActCa^+2^ amplitude during seizure onset and ictal in OHSC. (**H**) Beeswarm plot of time-averaged mean normalized ActCa^+2^ amplitude during seizure onset and ictal activity in vivo.

During seizure, the calcium amplitude of individual neurons also increased for the active state (Figure 5B and D). The baseline active calcium amplitude (ActCa^+2^_t=0_) was calculated for each neuron by averaging all active state values during the baseline time period and used to compute normalized 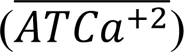. On average, the fold increase of ATCa^+2^ during seizure onset was 1.08 ± 0.006 in OHSC (n = 82 seizures; p < 0.001 paired t-test), and 1.20 ± 0.13 in vivo (n = 5 seizures; p = 0.09) (Figure 5G-H). During frank seizure, the average fold increase in active calcium amplitude was 1.37 ± 0.02 in OHSC (p < 0.001), and 1.40 ± 0.17 in vivo (p = 0.04) (Figure 5G-H). The observed increase in calcium amplitude is present even when accounting for the slower decay kinetics of GCaMP (Figure S5 A-C). This reveals that neurons do not enter a static “active” state, but rather, that intra-cellular perturbations allow for higher levels of calcium signal to occur during seizure, perhaps through high frequency action potential firing (*26*), action potential broadening (*27*), and/or PDS (*28*).

### How synchrony and calcium amplitude changes contribute to total seizure activity

With synchrony and intra-cellular calcium amplitude precisely quantified on the individual neuron scale, the relative contribution of each source to population means can be calculated (Figure 6). The fraction of the increased population signal that comes from synchronous activity, as either recruitment or reactivation, is calculated compared to 0% activation. The fraction from active state and subthreshold amplitude changes is calculated compared to the same level of activity, with active state and subthreshold amplitudes equal to amplitudes calculated during inter-ictal, non-epileptiform activity (see Supplemental Methods for details). The total contribution from active state neurons is represented by the summation of the contributions from network synchrony and increased ActCa^+2^ amplitude. On average in OHSC, the contributions to the increased population signal during seizure onset are 9.0% ± 0.8 recruitment, 17.3% ± 0.1 re-activation, 4.3% ± 0.5 increase in ActCa^+2^ amplitude, and 69.4% ± 1.5 increase in STCa^+2^ amplitude (n = 82 seizures; Figure 6C). In vivo, the increase in the population mean during seizure onset is 0.2% ± 0.02 recruitment, 15.7% ± 3.5 re-activation, 17.2% ± 4.1 increase in ActCa^+2^ amplitude, and 66.9% ± 7.6 increase in STCa^+2^ amplitude (n = 5 seizures; Figure 6F). This substantiates a new finding that the majority of the signal increase during seizure onset is due to changes in the amplitude of intra-cellular calcium in neurons while they are in the subthreshold calcium state. In other words, there are widespread, low-level calcium increases during seizure onset.

**Fig. 6.**
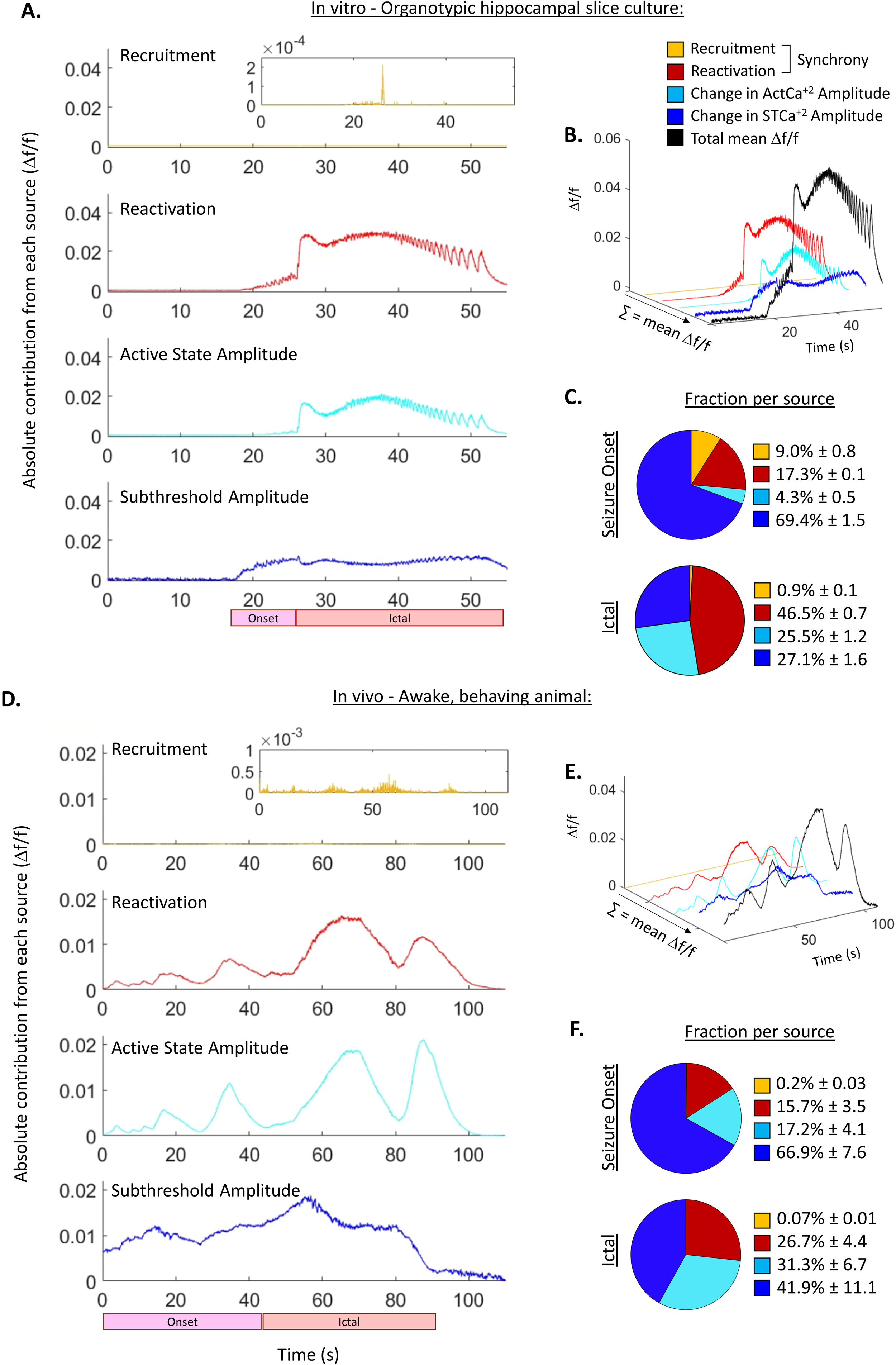
Quantifying sources of increased population calcium signal during seizure. (**A**) Plots of the absolute contributions of recruitment, reactivation, change in mean active calcium amplitude (*ActCa*^+2^), and mean subthreshold calcium amplitude (S*TCa*^+2^) in an example seizure recorded from an OHSC. (**B**) 3D line plot of the 4 sources of ictal signal and the total mean Δf/f signal in black from the example seizure in (A). The 4 sources (recruitment (yellow), reactivation (red), changing in ActCa^+2^ (cyan), and change in STCa^+2^ amplitude(blue)) summate into the population level mean Δf/f signal (black). (**C**) Pie chart of the mean relative contribution of each source during seizure onset and ictal activity, n = 82 seizures from OHSC. Relative contribution = absolute contribution/total Δf/f signal. (**D**) Plots of the absolute contributions of recruitment, reactivation, change in mean active calcium amplitude (*ActCa*^+2^), and mean subthreshold calcium amplitude (S*TCa*^+2^) in an example seizure recorded in vivo. (**E**) 3D line plot of the 4 sources of ictal signal and the total mean Δf/f signal in black from the example seizure in (D). The 4 sources (recruitment (yellow), reactivation (red), changing in ActCa^+2^ (cyan), and change in STCa^+2^ amplitude(blue)) summate into the population level mean Δf/f signal (black). (**F**) Pie chart of the mean relative contribution of each source during seizure onset and ictal activity, n = 5 in vivo seizures. Relative contribution = absolute contribution/total Δf/f signal.

During frank seizure in OHSC, the population signal arises from: 0.9% ± 0.1 recruitment, 46.5% ± 0.7 re-activation, 25.5% ± 1.2 ActCa^+2^ amplitude, and 27.1% ± 1.6 STCa^+2^ amplitude (Figure 6C). In vivo, the ictal population signal is 0.07% ± 0.01 recruitment, 26.7% ± 4.4 re-activation, 31.3% ± 6.7 increased ActCa^+2^ amplitude, and 41.9% ± 11.1 increased STCa^+2^ amplitude (Figure 6F). While subthreshold amplitude changes are still a significant factor during frank seizure, the relative contribution from active state neurons increases during ictal activity (Figure S6), with high levels of sustained activity in previously recruited cells. The main findings are robust to different experimental conditions, including version of GCaMP (Figure S7A), chronic versus periodic sampling (Figure S7B-C), method of ROI selection (Figure S7B-C), or spatial down-sampling (Figure S8). Quantifying the ictal contribution of subthreshold neurons appears to be the most susceptible to different experimental conditions and underscores the necessity of optimizing imaging conditions and ROI selection methods to capture all of the relevant neuronal activities.

## DISCUSSION

The quantifiable clinical hallmark of an epileptic seizure is manifest as an increase in the amplitude of EEG. Many prior studies have evaluated the intracellular correlates of ictal activity in vivo (*9, 29*) and in vitro (*30, 31*), but there remains a gap in understanding how intracellular activity of individual neurons contributes to population-level extracellular signals. Using simultaneously recorded somatic calcium transients and local field potentials, we found that cell intrinsic changes (i.e. the amplitude increases in individual neurons) were the largest contributor to the increased population signal during seizure onset. The high spatial resolution of calcium imaging also allowed us to resolve neuronal recruitment dynamics, substantiating the new finding that recruitment is a minor contributor to total network synchrony during seizure.

We were surprised to find that the change in subthreshold calcium activity was the single largest contributor to population activity during seizure onset. One way in which elevated subthreshold calcium amplitude could be maintained is through the combination of increased excitatory and inhibitory drive (allowing EPSP-mediated calcium influx that does not induce neuronal firing (*32*). Low frequency action potential firing may also occur in the subthreshold state and contribute to the observed increases in subthreshold calcium; however, it is unlikely that high frequency firing would remain subthreshold. The increased amplitude of subthreshold calcium persists during frank seizure, but the relative contribution of subthreshold calcium decreased as ictal activity progressed, consistent with the gradual collapse of surround inhibition associated with seizure propagation (*11, 33, 34*), (a requisite for seizure onset in our prior modeling study in this preparation (*35*)). Subthreshold calcium may also be increased by alterations to low-voltage activated channels (such as the T-type VGCC (*36*)), or changes in extracellular potassium. The combined contribution of an increased number of active state neurons and the increased amplitude of activity in those cells revealed that the majority of the ictal signal is generated by active state neurons, both in vitro and in vivo.

This study takes advantage of unique properties of calcium imaging to address questions that microelectrodes (MEA) cannot. The extracellular field recording largely comes from postsynaptic potentials (PSPs) (*37*). Since PSPs are inherently slower than action potentials, and arrive asynchronously at different locations, the extracellular electric field they generate is temporally broad, thus summating more readily than the brief extracellular voltage transients associated with action potentials. Calcium imaging is an imperfect proxy for neuronal electrical activity but is well-suited for capturing the effects of the PSP (Figure 1 and 2). Intracellular calcium transients comprise a lowpass filtered version of neuronal spiking (similar to field potential). Thus, calcium imaging provides an experimentally accessible mean by which to sum the activity of individual members of a seizing neural population.

Recent studies have begun to use new spike sorting techniques to investigate the activity of individual neurons during seizures recorded with MEAs in human epilepsy (*7, 38*). Modeling studies of the population response generated by a single neuron spiking have also attempted to bridge the gap between neuronal spiking activity and population activity measures (*39–41*). Both of these approaches begin from the identified spiking of a single cell. Our findings suggest that this spiking-centric approach may miss key features of the ictal signal. The widespread increase in subthreshold calcium suggests that during seizure onset a large population of neurons is sitting closer to action potential firing threshold. From this network state of high subthreshold calcium, it may be possible to trigger synchronized, ictal activity in a variety of ways. This is in line with our previous finding that seizures do not initiate in repeatable neuronal sequences (*20*) and suggests that many neurons would be able to drive ictal responses once the network has transitioned into a state of high subthreshold calcium. When interpreting MEA spiking data during seizure activity, it will thus be important to consider that the altered *response* to the spiking may be a critical factor in delineating the transition to seizure.

Future studies will be needed to assess the mechanisms responsible for the changes to network synchrony, and the active and subthreshold calcium states. Many studies have investigated pathological synchrony during seizure, with a myriad of potential mechanisms being identified (*42–44*). Despite these efforts, no consensus has been reached on how epileptic networks synchronize, illustrating the complexity of this process. Understanding the observed intra-cellular amplitude changes may offer new insights into seizure mechanism. Of particular interest, will be understanding the alterations in the subthreshold calcium state that occur during seizure onset. Uncovering these complex interactions will provide new insights into seizure initiation and potentially suggest new targets for treating seizures.

## Data availability

Data is available upon reasonable request to the corresponding author.

## Author contributions

LAL, KPL and KJS conceived this study. LAL and ZZ acquired data. LAL analyzed data. SNG, KPL and KJS obtained funding. LAL, KPL and KJS wrote the manuscript. All authors read, edited, and approved the final manuscript.

## Ethical Publication Statement

We confirm that we have read the Journal’s position on issues involved in ethical publication and affirm that this report is consistent with those guidelines.

