## Supplemental Text and Figures for "Cellular contributions to ictal population signals"

### SUPPLEMENTAL METHODS

#### Study design

As this is a study of ictal signals, and no treatments or modifications were performed, randomization and blinding are not applicable. For the same reason, there was no control group. Rather, single cell seizure dynamics were compared to periods without epileptiform activity in the same preparation. 8 in vivo seizures were captured from one APP/PS1 mouse (male, aged 5-months) with non-convulsive seizures. 5 seizures were analyzed as Type I (Figures 2 and 4-6) and 3 as Type II (Figure S1). Capturing neuronal resolution imaging data of spontaneous seizures in vivo is extremely rare. Many in vivo calcium imaging studies have relied on acute chemo-convulsant provoked seizures in non-epileptic brains (*1-4*). To date, only a single, partial spontaneous seizure has been reported with in vivo calcium imaging (and was of insufficient resolution for single cell analysis) (*5*). In comparison, spontaneous seizure-like events (referred to here as seizures) that occur in OHSCs are frequent and highly amenable to imaging. As described previously (*6*), our in vitro imaging paradigm was designed to capture approximately 1 seizure per day from each OHSC. Imaging began immediately after slice preparation and continued for 3 weeks. 82 seizures were captured from 10 OHSC, from 4 animals. No recorded seizures were excluded. In OHSC, only the first 10 seconds of ictal activity was analyzed many recordings terminated before the end of the seizure. In vivo, the imaging paradigm was longer and as all seizure terminations were captured, the entire ictal period was analyzed. Based on previous work which shows that seizure onset type evolves over time and that single cell sequences are stochastic during seizure recruitment (*6*), we treated each seizure as an independent event.

**In vivo virus injection**

Virus injection was performed as previously described (*7, 8*). The adult APP/PS1 mouse was anesthetized and head-fixed on a stereotaxic frame. A small hole overlying hippocampal CA1 at anterior-posterior (AP) -2.10 mm and medial-lateral (ML) 1.65 mm was drilled to expose the brain surface, and 1 μl of AAV1-hSyn-GCaMP6f virus solution (with a titer of ~ 1 $x10^{13}$/ml) was injected (dorsal-ventral (DV) -1.40 mm). After surgical closure, the animal was returned to the housing facility and was allowed to recover for 2 weeks.

**Endoscope and GRIN lens implantation surgery**

Surgery was performed as previously described (*7, 8*). The animal was anesthetized under isoflurane, head-fixed on a stereotaxic frame and the underlying soft tissue was cleared to expose the skull. The pial surface was exposed at two sites, one for implantation of a lens for calcium imaging (AP -2.10 mm, ML 1.50 mm), the other for insertion of a closely apposed electrophysiology probe (AP -2.80 mm, ML 2.60 mm). To prepare for lens implantation, the cortex above dorsal hippocampal CA1 was carefully aspirated with a vacuum needle. A gradient-index (GRIN) lens 4mm in length and 1mm in diameter (Inscopix) was implanted vertically just above CA1 (DV -1.2 mm from brain surface). A custom-made adjustable electrophysiology probe consisting of 7 stereotrodes was implanted (AP −2.8 mm, ML 2.6 mm from Bregma, −36 degrees relative to the AP axis, −39 degrees relative to the inter-aural axis; DV −1.7 mm from brain surface). A cerebellar screw was placed for use as ground. After closure, the animal was returned to the housing facility and allowed to recover. Three weeks later a microscope baseplate (Inscopix) was attached to the skull above the implanted GRIN lens.

### In vivo calcium imaging and electrophysiology recordings

For data acquisition, the miniscope (Inscopix) was mounted onto the baseplate, and a headstage pre-amplifier (Neuralynx) was connected to the electrophysiology probe. After a 30-minute habituation period, simultaneous calcium imaging (20 Hz) and electrophysiology recordings (local field potential recordings, 2 kHz sampling; multiunit recordings 32 kHz; filtered at 0.5–900 Hz) from each intracranial electrode were acquired, as previously described (*7*, *8*). Three 10-minute-long imaging sessions (with 5-minute off periods between to prevent photobleaching) were acquired with the animal in a state of quiet wakefulness. The electrophysiology signal was acquired continuously throughout the duration of the experiment.

### Organotypic Hippocampal Slice Culture (OHSC) preparation

Newborn DLX-cre mouse pups (Post-natal day (P) 1-2) received intracerebroventicular (ICV) injections of AAV-hSyn-GCaMP7f. Organotypic slice cultures were prepared from P6-P8 DLX-cre mice of either sex using the membrane insert technique (*9*), as described previously (*6*). Briefly, isolated hippocampi were cut into 400 μm slices on a McIlwain tissue chopper (Mickle Laboratory Engineering) and transferred to membrane inserts (Millipore) in six-well plates. Cultures were incubated in a custom-designed imaging system at 5% CO2, 36°C in 1 mL of Neurobasal-A growth medium supplemented with 2% B27, 500 μM GlutaMAX, and 0.03 mg/ml gentamycin (all from Invitrogen). Slices were maintained in the custom imaging system (see below).

For whole cell patch clamp experiments (Figure 1 and S1), OHSCs were prepared with the rocking coverslip method, as previously described (*10*). As above, OHSCs were prepared from P6-P8 DLX-cre mice crossed with Ai14 mice (which express tdtomato fluorescence following Cre-mediated recombination) of either sex. Slices were placed on poly-L-lysine coated glass cover slips (Electron Microscopy Sciences) in a 6-well tissue culture plate, with AAV9-hSyn-soma-GCaMP8m (10 μl per mL) included in the cell culture media. Slice cultures were maintained at 37°C, in 5% $CO_2$ on a rocking platform. Cell culture media was changed twice a week (without further inclusion of AAV9-hSyn-soma-GCaMP8m).

**OHSC chronic calcium imaging and local field potential recording**

On the day of slice culture preparation, the 6-well plate containing membrane slices was transferred to the custom imaging system, an inverted microscope constructed inside of a $CO_2$ incubator, as described previously (*6*, *11*). Slices were imaged every 6 hours to obtain an 85-second time series of green fluorescence at 35 Hz to record the spatiotemporal dynamics of calcium activity.

In some experiments, imaging was paired with electrical recordings of the extracellular field potential (*6*). Slice cultures were grown directly on top of the tungsten recording electrode. Data was collected at 1 KHz using a DC amplifier (Warner Instruments) with a 0.1 KHz filter.

**OHSC acute two-photon calcium imaging and whole cell recordings**

In a separate set of experiments, simultaneous two-photon calcium imaging and whole cell current clamp recordings were performed. For imaging/patching experiments, individual coverslips with OHSCs were transferred to the submersion chamber of a two-photon microscope between DIV 10-16. OHSCs were allowed to recover for approximately 15 minutes being perfused in cell culture media (at 1 ml/min), bubbled with 95%$O_2$/5% $CO_2$ at 33°C. Following recovery in cell culture media, perfusion was switched to aCSF (at 1ml/min, bubbled with 95%$O_2$/5% $CO_2$ at 33°C), containing (in mM) 124 NaCl, 1.25 NaH2PO4, 2.5 KCl, 26 NaHCO3, 2 CaCl2, 2 MgSO4, and 20 d-glucose.

Electrodes were pulled from borosilicate glass capillaries using a micropipette puller (Sutter Instruments) with resistance of 5–7 MΩ, filled with internal solution containing the following (in mm): 135 K+-gluconate, 5 NaCl, 2 $MgCl_2$, 10 HEPES, 0.4 $Na_3GTP$, 4 $Na_2ATP$, with osmolarity 290 mOSm, pH 7.25-7.35, adjusted with KOH. To aid in visualizing the pipette under fluorescence microscopy, 0.2% of 1mg/mL of Alexa Fluor 594 biocytin was added to the internal solution. Patching was done under 920 nm excitation, with a green 525/50 emission filter (Thor labs) to locate GCaMP-positive neurons. Interneurons were identifiable by DLX-driven expression of tdtomato. Resting membrane potential (RMP) was measured after whole-cell configuration was reached. Membrane potential was corrected for liquid junction potential of -14.3 mV. RMP of analyzed neurons was between -72 and -76 mV. All neurons that were recorded during spontaneous seizure activity were included (n = 3 pyramidal cells and n = 3 interneurons from the CA1 region, from 6 OHSCs).

Current clamp recordings were acquired with a Multiclamp amplifier (Multiclamp 700B, Molecular Devices) with Clampex10 software (Molecular Devices). Signals were sampled at 10 KHz and filtered at 2 KHz. Recordings were made in gap-free mode to capture spontaneous action potentials and postsynaptic potentials. Simultaneous 30 Hz, two-photon calcium imaging was performed, with a 16x, 0.8N.A. water-immersion objective (Nikon), on a custom-built two-photon microscope (*12*) equipped with a Mai Tai 80MHz Ti:sapphire laser (Spectra-Physics), and a resonant-galvo-galvo scanhead (Vidrio RMR). GCaMP calcium signal was excited at 920 nm, with a bandpass emission filter at 525/50 (Chroma). Image acquisition was performed by ScanImage software (Vidrio Technologies) (*13*). Both modalities were synchronized by triggering the ScanImage acquisition with Clampex. Analysis of current clamp recordings was done with Clampex11 software and Mini Analysis Software.

**Pre-processing of calcium signal**

*In vivo*: Neuronal regions of interest (ROI) were identified with the Fiji plugin TrackMate (*14*), using a Laplacian of Gaussian filter to identify GCaMP-positive neuronal cell bodies. TrackMate was used to follow the position of the same ROI over time and correct for motion artifact. The mean pixel intensity was extracted from the somatic ROI and a surrounding anulus ROI of neuropil. The calcium traces were normalized as Δf/f, and corrected for neuropil contamination as:

$$\frac{\Delta f}{f} corrected = \frac{\Delta f}{f} soma - \left(0.95 * \frac{\Delta f}{f} neuropil\right)$$

The weighting factor 0.95 was determined empirically to maximize the subtraction while avoiding over-correction.

*OHSC*: GCaMP-positive cells were identified from a standard deviation projection (STD) of the high-resolution calcium recordings, which highlights pixels with dynamic signal. The Fiji plugin TrackMate (*14*) was used to automatically select ROIs. Somatic signal was corrected for neuropil contamination as:

$$F_{corrected} = F_{soma} - \left(0.8 * F_{neuropil}\right)$$

As above, the weighing factor 0.8 was determined empirically. F corrected was normalized as Δf/f, as described previously (*6*).

In a subset of neurons, the somatic and neuropil ROIs were hand drawn (Supplemental Figure 2A) to allow for high fidelity neuropil contamination correction:

$$F_{corrected} = F_{soma} - F_{neuropil}$$

In this case nearly complete neuropil subtraction can be performed, with potential contamination coming only from neuropil directly above/below the soma.

For processing of the calcium signal obtained during acute two-photon imaging and whole cell patch clamp, the somatic and surrounding neuropil ROIs were hand-drawn to isolate the calcium signal from the patched neuron. In this case, a weighing factor of 0.5 was determined to best remove neuropil contamination.

To account for the slower decay kinetics of GCaMP, in a subset of data, a deconvolution was performed with the subtraction of an exponential decay based on the kinetics of GCaMP7f (*16*) (Figure S5).

**Separation of subthreshold and active calcium states**

*In vivo:* We identified a window of baseline activity (i.e. without seizure or epileptiform activity) on a cell-by-cell basis using the absolute value of calcium traces high-pass filtered at 0.2 Hz to isolate periods without high-amplitude calcium transients. Active state detection threshold was calculated as median + 2.5 standard deviations from the entire baseline period for each neuron. This strategy was selected by optimizing the detection of high-amplitude calcium transients (while minimizing false positives) in a sub-population of randomly selected cells. The detection thresholding process was then applied to the entire neuronal population in an automated fashion.

*OHSC*: A baseline window (without seizure or epileptiform activity) of approximately 3 seconds was identified in each seizure recording from the mean Δf/f trace. OHSC have inter-ictal periods of relative quiescence in all neurons, thus the same baseline window was used to calculate threshold in all neurons. Threshold was set as mean + 3 standard deviations on a cell-by-cell basis.

The same detection thresholding method was used for data collected from the chronic 1-photon imaging and acute 2-photon imaging.

**Quantification of calcium sources**

The mean population calcium signal (mean Δf/f) can be calculated from 3 variables (Figure S6): the fraction of synchronously active neurons ($S = \frac{\#\left(ActCa^{+2}\right)}{total\,\#\,(neurons)}$), the mean active state calcium amplitude ($ActCa^{+2}$), and the mean subthreshold calcium amplitude ($STCa^{+2}$) as:

$$\frac{\Delta f}{f} = \left(ActCa^{+2} * S\right) + \left(STCa^{+2} * (1 - S)\right)$$

The first term represents the total contribution from active state neurons and the second from subthreshold neurons. The relative contribution of active state and subthreshold neurons can be found by diving by the total Δf/f signal (Figure S6).

We can also calculate the fractional increase from four sources during seizure. In this case, seizure (either seizure onset or frank seizure) is being compared to a reference period of physiological activity. In vivo, this was a period of quiet wakefulness without any epileptiform activity. In vitro, this was a period of inter-ictal activity. The reference period is defined here as t = 0. The exact duration of t = 0 varied depending on how much physiological activity was captured in the same recording as a seizure but was generally tens of seconds. The mean amplitude of $ActCa^{+2}$ and $STCa^{+2}$ was calculated from this period, as $ActCa^{+2}_{t=0}$ and $STCa^{+2}_{t=0}$. The contribution to the population Δf/f signal from network synchrony can then be calculated as:

Synchrony: $\left(ActCa^{+2}{}_{t=0} * S\right) + \left(STCa^{+2}{}_{t=0} * (1 - S)\right) - STCa^{+2}{}_{t=0}$

In other words, this is the contribution from an increasing number of active cells if the $ActCa^{+2}$ and $STCa^{+2}$ amplitude was unchanged from baseline. Network synchrony includes both recruitment and re-activation. The fraction of synchronous activity from each source can be isolated as:

Recruitment: $$\frac{ActCa^{+2}t=0 * \frac{\#\left(ActCa^{+2}recruited\right)}{total\#\ (\ neurons)}}{\left(ActCa^{+2}t=0 * \ S\right)}$$

Reactivation: $$\frac{ActCa^{+2}t=0 * \frac{\#\left(ActCa^{+2}reactivated\right)}{total\#\ (\ neurons)}}{\left(ActCa^{+2}t=0 * \ S\right)}$$

The fraction of the population signal which comes from $ActCa^{+2}$ and $STCa^{+2}$ amplitude changes can be calculated as:

Active calcium amplitude: $\left(ActCa^{+2} * \ S\right) - \left(ActCa^{+2}{}_{t=0} * \ S\right)$

Subthreshold calcium amplitude: $\left(STCa^{+2} * \ (1 - S)\right) - \left(STCa^{+2}{}_{t=0} * \ (1 - S)\right)$

In other words, the fractional increase is calculated as the difference in the total contribution from all neurons in the active state (or subthreshold state) and the contribution from the same number of neurons if the amplitude was the same as in the t=0 reference period.

**Varying Experimental Conditions**

In a subset of experiments (Figure S7), OHSCs were prepared from the same animal using the membrane insert technique as above but transfected with different versions of GCaMP. OHSCs were incubated in a 6-well tissue culture plate, with either AA9-hSyn-GCaMP6f, AAV9-hSyn-GCaMP7f or AAV9-hSyn-soma-GCaMP8m (5 μl per mL) included in the cell culture media. Cell culture media was changed twice a week (without further inclusion of AAVs). Between DIV8-12, the 6-well plate was transferred to the custom imaging system as above and were imaged 4 times per day to obtain an 85-second time series of green fluorescence at 35 Hz to record the spatiotemporal dynamics of calcium activity. Between imaging sessions, the OHSCs were returned to a standard $CO_2$ incubator. Under these conditions, neuronal ROI selection was found to be optimized by generating multiple standard deviation projections (STD) and then performing a simple ratio normalization (ROI method #2, Figure S7B-C) to use in ROI selection by TrackMate. ROI selection based on a single STD projection (ROI method #1) may bias data to preferentially select neurons active during the seizure. Under chronic imaging conditions, the difference between the ROI selection method was less apparent (Figure S7B-C).

**Statistics**

All data are reported as mean ± SEM. Tests of correlation were performed and reported as the Pearson correlation coefficient (r). Paired student *t*-tests were performed to test for statistical significance, as noted in text. In all cases, single neuron data was being compared to data from a reference period in the same cell. To test for effects of experimental conditions on the distribution of the fraction of signal from each source, 2-way ANOVAs were performed. Statistical analyses were performed with Microsoft Excel or Matlab (MathWorks).

1. M. Wenzel, J. P. Hamm, D. S. Peterka, R. Yuste, Acute Focal Seizures Start As Local Synchronizations of Neuronal Ensembles. *J Neurosci* **39**, 8562-8575 (2019).
2. X. Zhang, Z. Qiao, N. Liu, L. Gao, L. Wei, A. Liu, Z. Ma, F. Wang, S. Hou, J. Li, H. Shen, Stereotypical patterns of epileptiform calcium signal in hippocampal CA1, CA3, dentate gyrus and entorhinal cortex in freely moving mice. *Sci Rep* **9**, 4518 (2019).
3. T. K. Berdyyeva, E. P. Frady, J. J. Nassi, L. Aluisio, Y. Cherkas, S. Otte, R. M. Wyatt, C. Dugovic, K. K. Ghosh, M. J. Schnitzer, T. Lovenberg, P. Bonaventure, Direct Imaging of Hippocampal Epileptiform Calcium Motifs Following Kainic Acid Administration in Freely Behaving Mice. *Frontiers in neuroscience* **10**, 53 (2016).
4. M. Wenzel, J. P. Hamm, D. S. Peterka, R. Yuste, Reliable and Elastic Propagation of Cortical Seizures In Vivo. *Cell Rep* **19**, 2681-2693 (2017).
5. S. F. Muldoon, V. Villette, T. Tressard, A. Malvache, S. Reichinnek, F. Bartolomei, R. Cossart, GABAergic inhibition shapes interictal dynamics in awake epileptic mice. *Brain* **138**, 2875-2890 (2015).
6. L. A. Lau, K. J. Staley, K. P. Lillis, In vitro ictogenesis is stochastic at the single neuron level. *Brain*, (2021).
7. H. Zhou, K. R. Neville, N. Goldstein, S. Kabu, N. Kausar, R. Ye, T. T. Nguyen, N. Gelwan, B. T. Hyman, S. N. Gomperts, Cholinergic modulation of hippocampal calcium activity across the sleep-wake cycle. *Elife* **8**, (2019).
8. H. Zhou, H. Li, N. Gowravaram, M. Quan, N. Kausar, S. N. Gomperts, Disruption of hippocampal neuronal circuit function depends upon behavioral state in the APP/PS1 mouse model of Alzheimer's disease. *Sci Rep* **12**, 21022 (2022).
9. L. Stoppini, P. A. Buchs, D. Muller, A simple method for organotypic cultures of nervous tissue. *J Neurosci Methods* **37**, 173-182 (1991).
10. V. I. Dzhala, K. J. Staley, KCC2 Chloride Transport Contributes to the Termination of Ictal Epileptiform Activity. *eNeuro* **8**, (2021).
11. T. Jacob, K. P. Lillis, Z. Wang, W. Swiercz, N. Rahmati, K. J. Staley, A Proposed Mechanism for Spontaneous Transitions between Interictal and Ictal Activity. *J Neurosci* **39**, 557-575 (2019).
12. B. A. Costine-Bartell, L. Martinez-Ramirez, K. Normoyle, T. Stinson, K. J. Staley, K. P. Lillis, 2-Photon imaging of fluorescent proteins in living swine. *Sci Rep* **13**, 14158 (2023).
13. T. A. Pologruto, B. L. Sabatini, K. Svoboda, ScanImage: flexible software for operating laser scanning microscopes. *Biomed Eng Online* **2**, 13 (2003).
14. J. Schindelin, I. Arganda-Carreras, E. Frise, V. Kaynig, M. Longair, T. Pietzsch, S. Preibisch, C. Rueden, S. Saalfeld, B. Schmid, J. Y. Tinevez, D. J. White, V. Hartenstein, K. Eliceiri, P. Tomancak, A. Cardona, Fiji: an open-source platform for biological-image analysis. *Nat Methods* **9**, 676-682 (2012).
15. T. W. Chen, T. J. Wardill, Y. Sun, S. R. Pulver, S. L. Renninger, A. Baohan, E. R. Schreiter, R. A. Kerr, M. B. Orger, V. Jayaraman, L. L. Looger, K. Svoboda, D. S. Kim, Ultrasensitive fluorescent proteins for imaging neuronal activity. *Nature* **499**, 295-300 (2013).
16. H. Dana, Y. Sun, B. Mohar, B. K. Hulse, A. M. Kerlin, J. P. Hasseman, G. Tsegaye, A. Tsang, A. Wong, R. Patel, J. J. Macklin, Y. Chen, A. Konnerth, V. Jayaraman, L. L. Looger, E. R. Schreiter, K. Svoboda, D. S. Kim, High-performance calcium sensors for imaging activity in neuronal populations and microcompartments. *Nature methods* **16**, 649-657 (2019).

**SUPPLEMENTAL FIGURES**

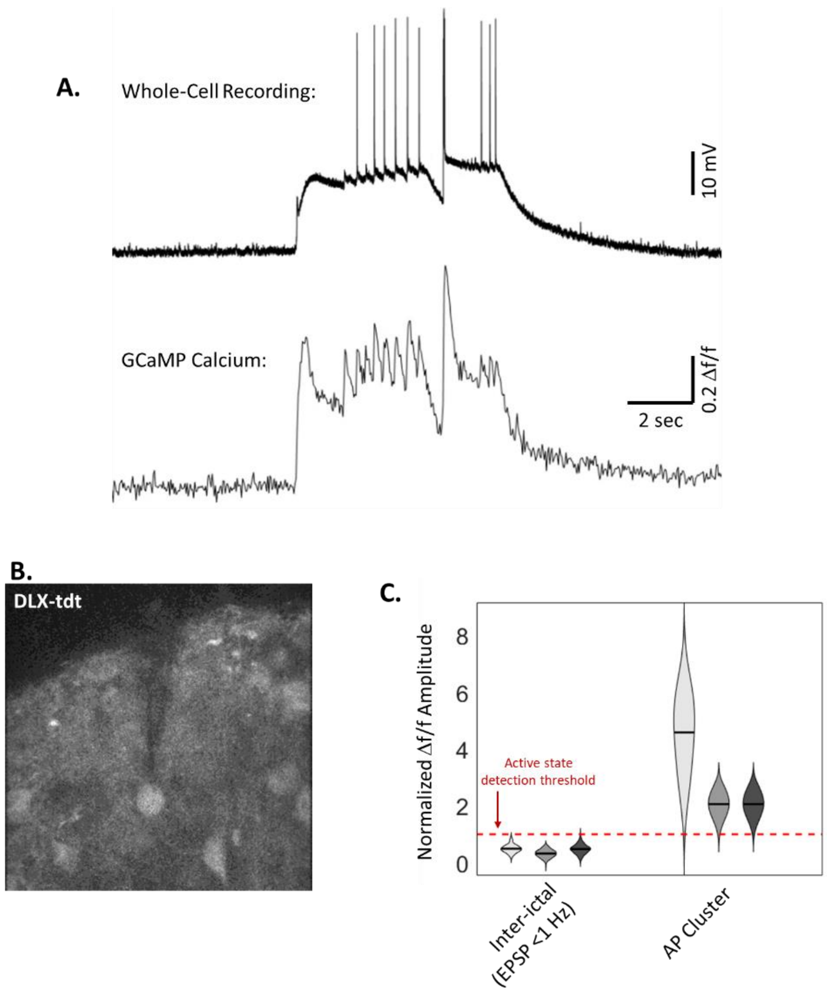


**Supplemental Fig. 1. Paired Calcium Imaging and Whole-cell Patch Clamp in Interneurons** (**A**) Paired current-clamp recording and soma-GCaMP8m calcium signal (Df/f) during spontaneous seizure. (**B**) DLX-tdt image showing patched interneuron. (**C**)Violin plot of calcium Δf/f amplitudes during inter-ictal activity and action potential clusters during spontaneous seizures, normalized to active state detection threshold per cell. (n = 3 interneurons, from 3 OHSCs).

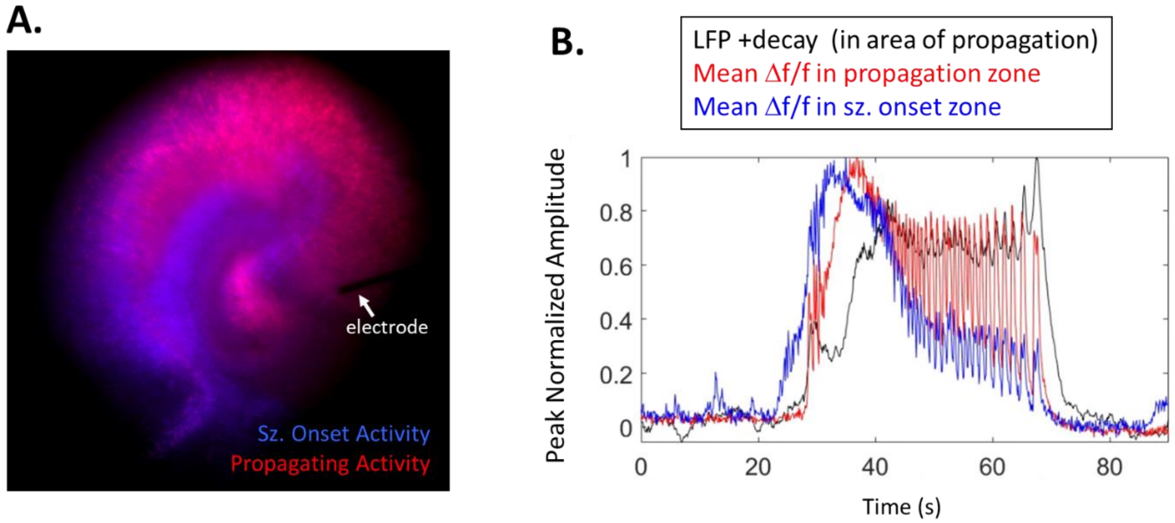


**Supplemental Fig. 2. Low correlation when electrode outside of seizure onset zone**

(**A**) Image of standard deviation projection of GCaMP7f calcium signal during seizure onset (blue) and propagating seizure activity (red) in an OHSC. (**B**) Mean Δf/f calcium signal of neurons within the seizure onset zone (blue), Mean Δf/f calcium signal of neurons in the area of seizure propagation, and the LFP signal from electrode in the propagation area (black; LFP was rectified, down-sampled and convolved with the decay kinetics of GCaMP7f, as in Figure 2). Note the appearance that the calcium signal precedes the electrical signal when the electrode is outside of the seizure onset zone.

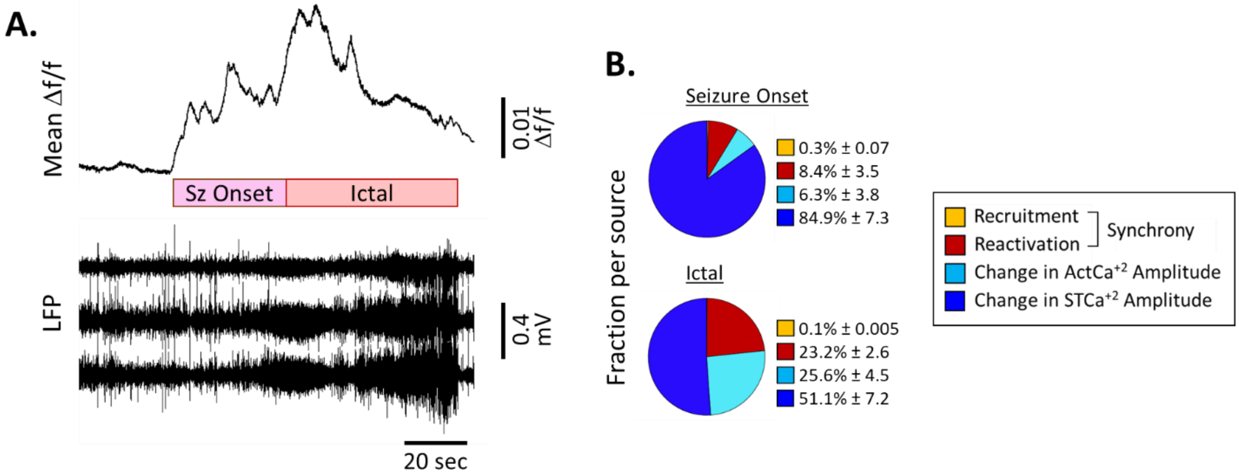


**Supplemental Fig. 3. Quantifying sources of increased population calcium signal during type II seizures**

(**A**) Example mean Δf/f calcium trace and paired local field potential (LFP) during a type II seizure recorded in vivo. Type II defined by high amplitude calcium activity which precedes increased electrical activity by several seconds. (**B**) Pie chart = mean percent of population increase from each source for type II seizures (recruitment (yellow), reactivation (red), change in active $Ca^{+2}$ amplitude (cyan), change in subthreshold $Ca^{+2}$ amplitude (blue)), n = 3 seizures.

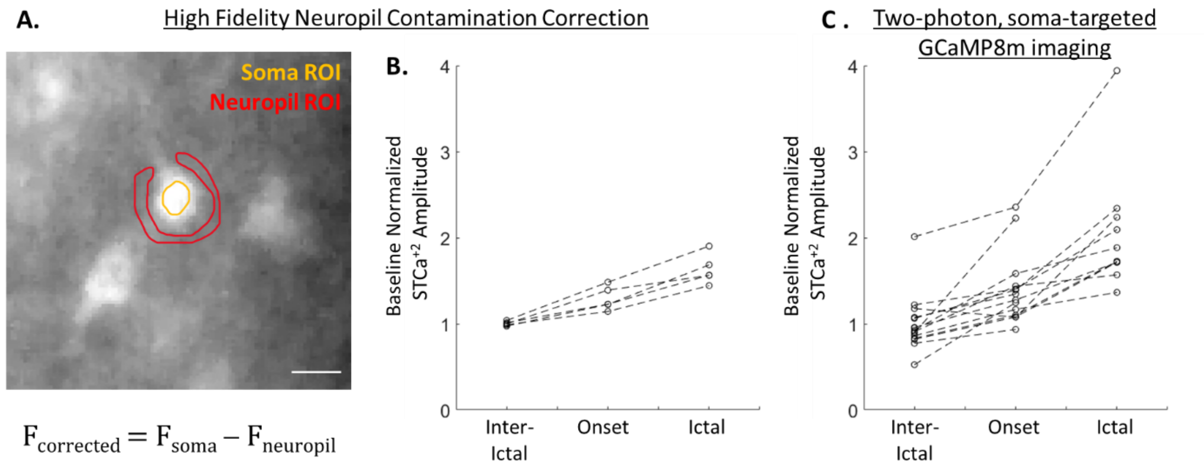


**Supplemental Fig. 4. Subthreshold amplitude changes robust to neuropil contamination corrections**

(**A**) Example of hand drawn soma and neuropil regions of interest in one-photon, GCaMP7f imaging. Scale bar = 10 µm. (**B**) Subthreshold calcium amplitude (normalized to baseline) during inter-ictal, onset and ictal periods with high fidelity neuropil contamination correction. n = 5 neurons; p = 0.003 (inter-ictal versus onset) and p < 0.001 (inter-ictal versus ictal), paired t-test. (**C**) Subthreshold calcium amplitude (normalized to baseline) during inter-ictal, onset and ictal periods in two-photon, soma-targeted GCaMP8m imaging. n = 15 neurons; p < 0.001 (inter-ictal versus onset) and p < 0.001 (inter-ictal versus ictal), paired t-test.

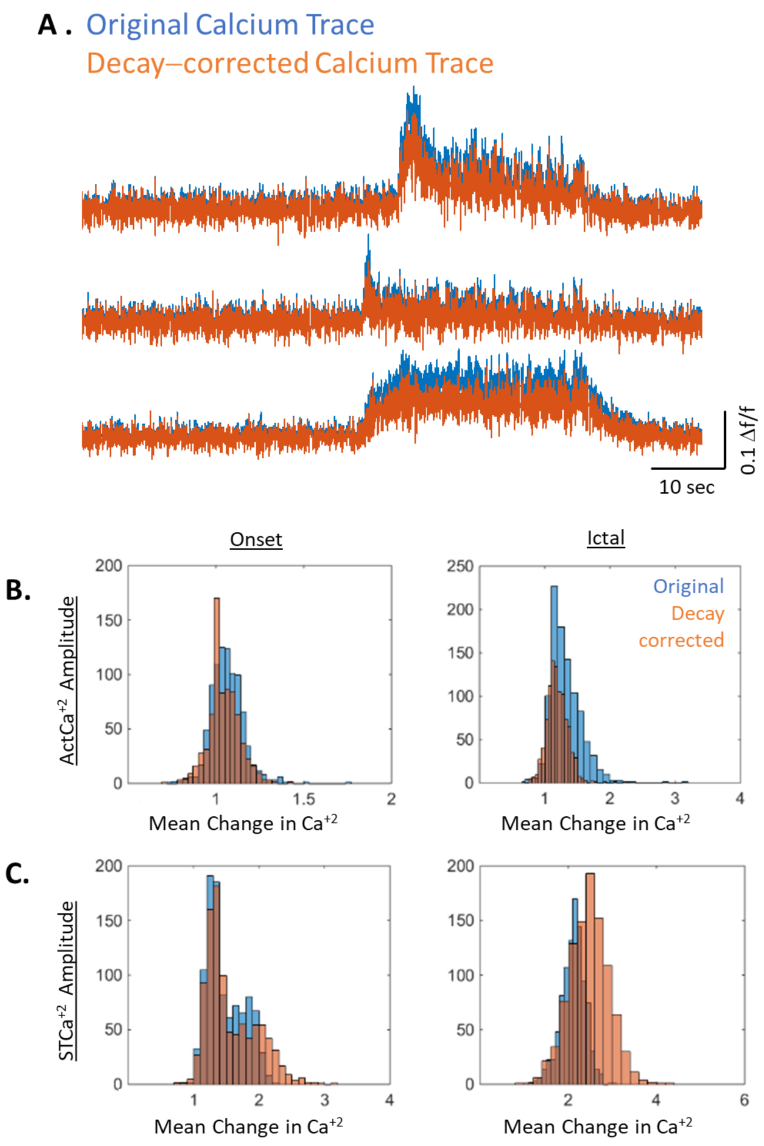


**Supplemental Fig. 5. Active state calcium amplitude increases robust to GCaMP decay kinetic correction**

(**A**) Example calcium traces from individual neurons (original: blue) and after subtraction of GCaMP7f decay kinetics (blue). (**B**) Histograms of the mean change in the active state calcium amplitude during seizure onset and ictal period from original (blue) and decay subtracted (orange) data (**C**) Histograms of the mean change in the subthreshold calcium amplitude during seizure onset and ictal period from original (blue) and decay subtracted (orange) data.

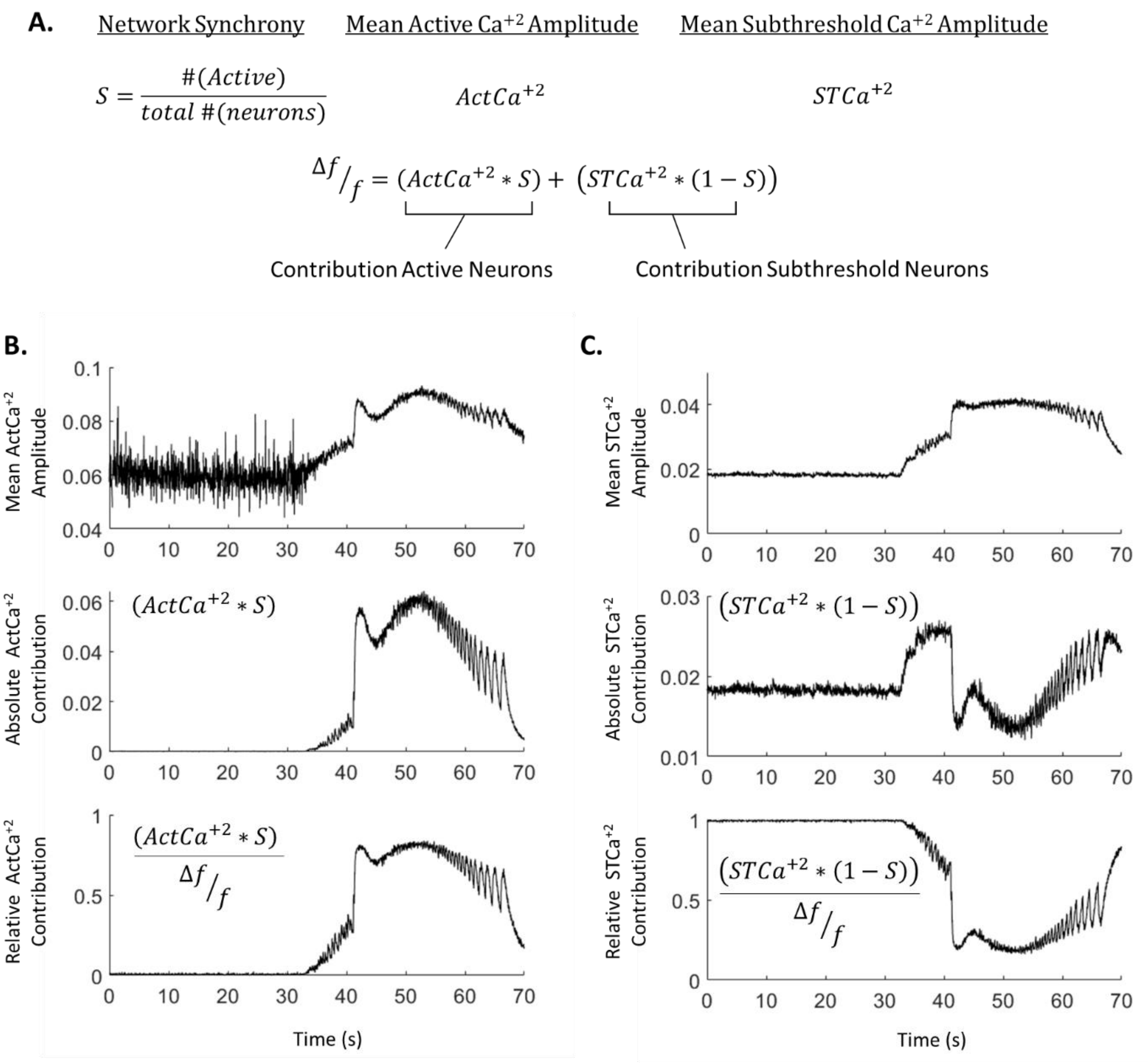


**Supplemental Fig. 6. Quantifying contribution from active and subthreshold neurons**

(**A**) Equation for quantifying the contribution from active neurons and subthreshold neurons, which can be calculated from 3 terms: network synchrony ($S = \frac{\#(ActCa^{+2})}{total\ \#(neurons)}$), mean active calcium amplitude ($ActCa^{+2}$), and mean subthreshold calcium amplitude ($STCa^{+2}$). (**B**) Example plot of the mean $ActCa^{+2}$ amplitude over time, absolute contribution of active state neurons over time ($ActCa^{+2}$ amplitude * fraction of active neurons), and relative contribution of active state neurons (($ActCa^{+2}$ amplitude * fraction of active neurons)/total Δf/f signal) over time from an example seizure in OHSC. (**C**) Example plot of the mean $STCa^{+2}$ amplitude over time, absolute contribution of subthreshold neurons over time ($STCa^{+2}$ amplitude * fraction of subthreshold neurons), and relative contribution of subthreshold neurons (($STCa^{+2}$ amplitude * fraction of subthreshold neurons)/total Δf/f signal) over time from an example seizure in OHSC.

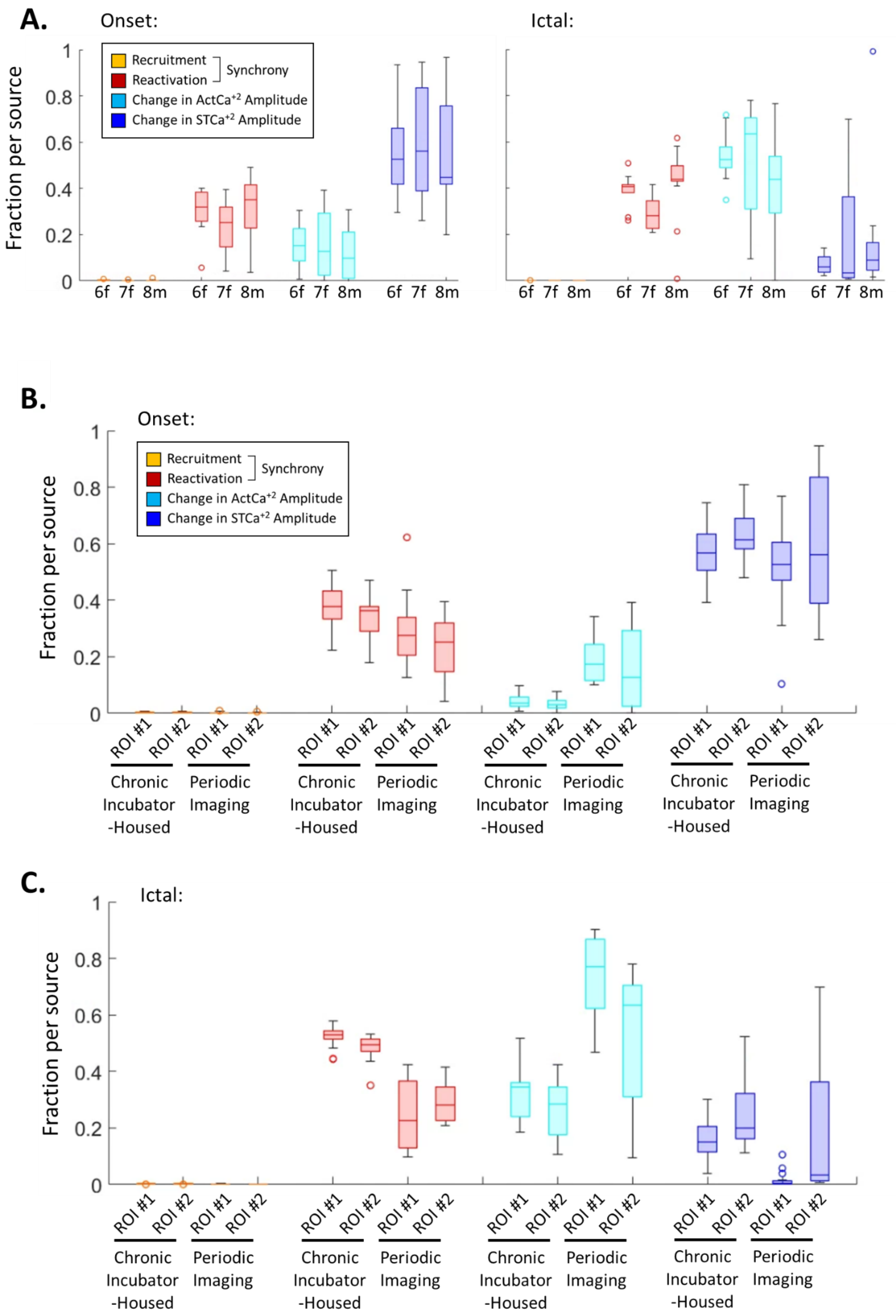

A.
Onset:
Ictal:
Fraction per source
Recruitment
Reactivation
Synchrony
Change in ActCa+2 Amplitude
Change in STCa+2 Amplitude
6f 7f 8m
B.
Onset:
C.
Ictal:
ROI #1 ROI #2
Chronic Incubator -Housed
Periodic Imaging

**Supplemental Fig. 7. Variability based on experimental conditions**

(**A**) Boxplot of the fraction of population signal from recruitment, reactivation, change in mean active calcium amplitude ($ActCa^{+2}$), and mean subthreshold calcium amplitude (S$TCa^{+2}$) during seizure onset (left) and ictal activity (right) in OHCS using either GCaMP6f, GCaMP7f or soma-targeted GCaMP8m. No significant interaction was found between GCaMP versions and fraction per source (2-way ANOVA), n = 11, 16 and 13 seizures from 4 OHSCs per condition. B) Boxplot of the fraction of population signal from recruitment, reactivation, change in mean active calcium amplitude ($ActCa^{+2}$), and mean subthreshold calcium amplitude (S$TCa^{+2}$) during seizure onset in either continuous incubator-housed imaging (n = 15 seizure, from 2 OHSC) or periodic imaging conditions (n= 16 seizures, from 2 OHSCs), using either raw amplitude ROI selection (ROI #1) or amplitude normalized ROI selection (ROI #2), all collected with GCaMP7f. C) Same as (B), but for ictal period.

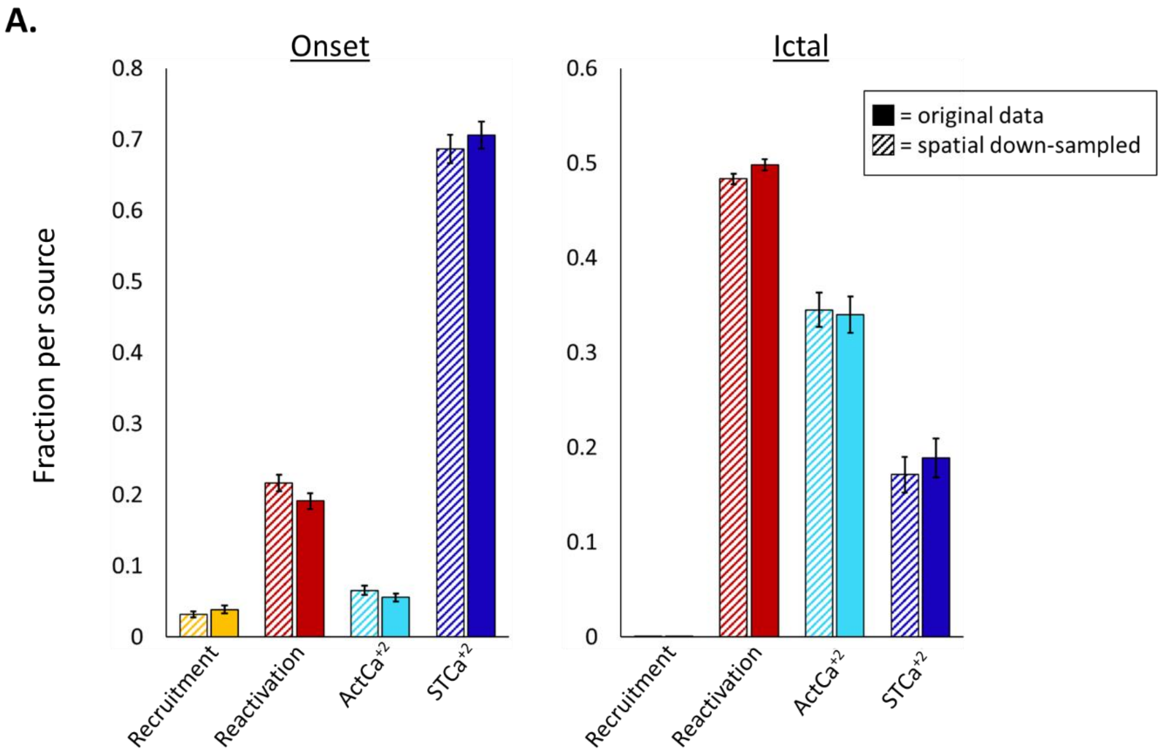


**Supplemental Fig. 8. Quantifying sources of increased population calcium signal after spatial subsampling**

(**A**) Mean percent of population increase from each source during seizure onset (recruitment, synchrony, change in active $Ca^{+2}$ amplitude, change in subthreshold $Ca^{+2}$ amplitude), n = 20 seizures in OHSC. Subsampling was performed by averaging the signal of neurons that fell within 14 μm of each other. (**B**) Mean percent of population increase from each source during ictal activity.
